## Supplementary material for "Lung spatial profiling reveals a T cell signature in COPD patients with fatal SARS-CoV-2 infection": Online supplement

**Methods**

**Study population**

The study included one main cohort of patients who died of COVID-19 infection, and a secondary cohort of uninfected subjects. The main cohort included seven never-smokers and eleven ever-smokers with and without COPD who died from COVID-19 pneumonia following SARS-CoV2 infection at the University of Navarra Hospital, Pamplona, Spain. The secondary cohort included 19 subjects who did not have COVID-19 infection and underwent lung volume reduction surgery or transplant for the treatment of severe emphysema, or lung resection for a solitary peripheral nodule (the lung tissue studied was at least 10 centimeters away from the nodule)^1^. In this second cohort, none of the subjects studied had evidence of respiratory tract infection at the time of lung tissue sampling, and COPD was diagnosed according to the Global Initiative for Obstructive Lung Disease^2^ recommendations.

All the study subjects defined as ever-smokers had a smoking history of at least 10 pack/years. All ever-smokers and COPD patients had ceased smoking for at least 1 year prior to the study.

**Spatial gene profiling of parenchyma, bronchiole, and vessel tissues**

The Nanostring GeoMX Digital Spatial Profiler (DSP) RNA slide preparation was followed according to the manufacturer’s instruction (<https://nanostring.com/products/geomx-digital-spatial-profiler/geomx-rna-assays>). Briefly, the slides were baked, deparaffinized, and antigen retrieval was performed using Tris-EDTA pH 9.0. Next, samples were digested with proteinase K and hybridization was performed overnight using the Cancer Transcriptome Atlas and the COVID Immune Response Atlas RNA detection probes (https://www.nanostring.com/products/geomx-digital-spatial-profiler/geomx-rna-assays/geomx-cancer-transcriptome-atlas/). The same sections were then stained with fluorescently labeled SYTO13, pan-cytokeratin, and Cluster of Differentiation (CD)45 to identify tissue regions of interest (ROIs, **Figures 2A**). For each lung section, we randomly selected 16 total regions of interest (ROI) from the parenchyma, airways and vessels, uniformly spread throughout the section to ensure sample randomization. Each ROI was subdivided into compartments based on fluorescent cell-specific markers, and serial UV illumination of each compartment was used to sequentially collect the probe barcodes from the different cell types as described elsewhere^3^. Once all of the indexing oligonucleotides were collected into a 96 well plate they were counted using next-generation sequencing. Each ROI was indexed using Illumina’s i5xi7 dual-indexing system and the cDNA libraries were pair-end sequenced on the NextSeq instrument (Illumina). Of the total of 1839 genes, 1803 were reliably measured across the dynamic range of gene expression panels (log2 normalized gene expression > 90th percentile of the expression of the negative control probes).

**Spatial protein profiling of parenchyma, airways, and vessel structures**

We used GeoMX Nanostring Digital Spatial Profiler to perform proteomic and transcriptomic analyses of the tissue sections from each subject. The samples were incubated with 41 oligo-labeled primary antibodies (See **Supplemental** **Table E2**): a Human Immune Cell Profiling Core, a Human Immune Activation Status Panel, a Human Immune Cell Typing Panel, a COVID-19 Immune Monitoring Panel, and custom-labelled antibodies against Syndecan-1, CD10, CD21 (all from abcam, Cambridge, MA). The slides were then loaded onto the GeoMX DSP and 16 ROIs were randomly selected per slide, and analyzed for protein expression. All the indexing oligonucleotides were collected into a 96 well plate, and were then hybridized to fluorescent barcodes using GeoMX Hyb Codes. After hybridization, samples were processed using the using the nCounter system according to the manufacturer’s instructions.

**ACE2 and CD45RO immunofluorescence staining of FFPE lung tissue sections**

The full staining protocol is described elsewhere^4^. Lung FFPE sections were deparaffinized, and antigen (Ag) retrieval was performed by treating the slides immersed in 0.01 M sodium citrate and 2 mM citrate buffer (pH 6.0) in a microwave. To identify ACE2- and CD45RO-positive cells within the alveolar and bronchiolar epithelium, lung sections were incubated with: 1) murine monoclonal antibody to CD45RO at 37°C for 2 hours (1:100, Abcam, Cambridge), followed by Alexa 488 conjugated goat anti-mouse IgG (H+L) at 37°C for an hour (1:100, Fisher Scientific); or 2) rabbit polyclonal to ACE2 (1:100, Abcam) at 37°C for an hour followed by Alexa-488 conjugated goat anti-rabbit IgG (H+L) at 37°C for an hour (1:100, Fisher Scientific, Hampton, NH). Then, a goat polyclonal antibody to EPCAM was added overnight at 4°C (1:100, Bio-Techne, Minneapolis, MN), followed by Alexa-568 conjugated rabbit anti-goat IgG (H+L) at 37°C for an hour (1:100, Fisher Scientific).

To identify ACE2- and CD45RO- positive cells in the endothelial wall, lung sections were incubated at 37°C for 2 hours with rabbit polyclonal antibody to ACE2 (1:50, abcam, ab15348), or murine monoclonal antibody to CD45RO (1:100, abcam,) at 37°C for an hour, followed by Alexa-488-conjugated goat anti-rabbit IgG (H+L) or anti-mouse IgG (1:100, Fisher Scientific), respectively. Then, a sheep polyclonal to Von Willebrand Factor was applied for 2 hours at 37°C (1:50, abcam), followed by Alexa 647 conjugated donkey anti-sheep IgG (H+L) at 37°C for an hour (1:100, abcam). Lung sections were also immuno-stained with appropriate isotype-matched non-immune control antibodies.

For each sample, 20 randomly-selected high-magnification fields for each tissue type (alveoli, airways, and vessels) were evaluated using an Olympus epi-fluorescence microscope (Olympus Global Corporation). The area of the alveolar, bronchiolar basement membrane (BM), and vascular tissue measured was quantified using a Metamorph software (Molecular Devices, San Jose, CA) as we previously published^4,5^. The number of parenchymal, vascular, and bronchial ACE2^+^ and CD45RO^+^ cells were counted in separate analysis by two independent investigators blindly, and normalized by each tissue area. The data were expressed as either mean or median number of parenchymal, bronchial, and vascular ACE2^+^ and CD45RO^+^ cells/ alveolar, bronchial, and vascular tissue areas, respectively.

**Statistics**

The cellular proportions of the immune cell types between tissue types and disease conditions were inferred using the R package “SpatialDecon”, a deconvolution method developed for the NanoString GeoMX RNA assay. The “Lung plus neutrophil” panel and the “Tcell_Adult_Lung_10x” panel were used to predict the cellular proportions of each ROI.

**Tables**

**Table E1. Demographic and clinical characteristics of patients without COVID-19**

|  | n | NS | ES | COPD | p |
| --- | --- | --- | --- | --- | --- |
| Total Partecipants (n) | 19 | 4 | 6 | 9 |  |
| Age | 19 | 63 [58-69] | 71 [67, 76] | 64 [62,68] | 0.199 |
| Gender (M/F) | 19 | 2/2 | 5/1 | 7/2 | 0.319 |
| Smoking habit (current/former smoker) | 19 | NA | 0/6 | 0/9 | NA |
| FEV1 (% predicted) | 18 | 91% (16) | 90% (18) | 38% (24) | 0.0004 |
| Comorbidities |  |  |  |  |  |
| Hypertension | 19 | 3 (75%) | 3 (50%) | 3 (30%) | 0.164 |
| Other cardiovascluar diseases | 19 | 2 (50%) | 2 (30%) | 4 (44%) | 0.669 |
| Diabetes Mellitus | 19 | 2 (50%) | 0 | 1 (11%) | 0.052 (NS vs ES) |
| Medications |  |  |  |  |  |
| Inhaled corticosteroids | 18 | 0 | 0 | 6 (67%) | 0.009 |
| LABA/SABA/LAMA | 18 | 0 | 0 | 8 (89%) | 0.002 |
| Oral corticosteroids | 18 | 0 | 1 (17%) | 3 (33%) | 0.187 |
| ACEi/ARB | 18 | 2 (50%) | 2 (30%) | 2 (22%) | 0.633 |

Data are expressed as the median [interquatile range] for continuous variables and n (%) for categorical variables. NS: never-smokers; ES: ever-smokers without COPD; LABA: long-acting beta-agonists; SABA: short-acting beta-agonists; LAMA: long-acting muscarinig agents; ACEi: ACE-inhibitors; ARB: angiotensin receptor blockers.

**Table E2: Nanostring GeoMX Protein Targets**

| Protein | Code Class | Protein Group |
| --- | --- | --- |
| S6 | Control | Housekeepers;All targets |
| Rb IgG | Negative | Background;All targets |
| Ki-67 | Endogenous | Proliferation;All targets |
| CD45 | Endogenous | Total Immune;All targets |
| PD-1 | Endogenous | T cells;Checkpoint;T cell Activation;All targets |
| CD68 | Endogenous | M2 Macrophage;Myeloid;Macrophage;All targets |
| GZMB | Endogenous | T cell Activation;Cytotoxicity;All targets |
| Ms IgG1 | Negative | Background;All targets |
| GAPDH | Control | Housekeepers;All targets |
| Histone H3 | Control | Housekeepers;All targets |
| CTLA4 | Endogenous | T cells;Checkpoint;T cell Activation;Th cells;All targets |
| PD-L1 | Endogenous | Checkpoint;Myeloid Activation;All targets |
| Fibronectin | Endogenous | Stroma;Fibroblasts;All targets |
| CD20 | Endogenous | B cells;All targets |
| CD4 | Endogenous | T cells;Myeloid;Th cells;All targets |
| CD8 | Endogenous | T cells;CD8 T cells;All targets |
| Ms IgG2a | Negative | Background;All targets |
| HLA-DR | Endogenous | Antigen Presentation;MHC2;All targets |
| CD3 | Endogenous | T cells;All targets |
| PanCk | Endogenous | Tumor;Epithelial;All targets |
| Beta-2-microglobulin | Endogenous | Tumor;Antigen Presentation;All targets |
| CD11c | Endogenous | DC;Myeloid;All targets |
| SMA | Endogenous | Stroma;All targets |
| CD56 | Endogenous | NK cells;All targets |
| SARS-CoV-2 Spike | Endogenous | Virus;All targets |
| Cathepsin L/V/K/H | Endogenous | Protease;All targets |
| TMPRSS2 | Endogenous | Protease;All targets |
| DDX5 | Endogenous | Immune Response;All targets |
| ACE2 | Endogenous | Viral Receptor;All targets |
| CD27 | Endogenous | T cells;T cell Activation;All targets |
| CD80 | Endogenous | Myeloid;Myeloid Activation;All targets |
| CD40 | Endogenous | Myeloid;Myeloid Activation;All targets |
| CD44 | Endogenous | T cell Activation;All targets |
| CD25 | Endogenous | T cells;T cell Activation;Tregs;All targets |
| PD-L2 | Endogenous | Checkpoint;All targets |
| CD127 | Endogenous | T cells;Naive and Memory;All targets |
| ICOS | Endogenous | T cell Activation;All targets |
| CD14 | Endogenous | Myeloid;Monocyte;All targets |
| CD45RO | Endogenous | T cells;Memory;All targets |
| FOXP3 | Endogenous | T cells;Th cells;Tregs;All targets |
| CD34 | Endogenous | Hematopoietic;All targets |
| FAP-alpha | Endogenous | Stroma;Fibroblasts;All targets |
| CD163 | Endogenous | M2 Macrophage;Myeloid;Macrophage;All targets |
| CD66b | Endogenous | Myeloid;Neutrophil;All targets |
| CD21 | Endogenous | Custom Target;All targets |
| Syndecan-1 | Endogenous | Custom Target;All targets |
| CD10 | Endogenous | Custom Target;All targets |

**Figure Legends**

**Figure E1. (A)** Thoracic computed tomography (CT) scans from never-smokers (NS), ever-smokers (ES), and patients with COPD within 4-10 days before death due to SARS-CoV-2 infection were assessed for emphysema (% lower attenuation area (LAA) <-950 Hounsfield units (HU), using Vida Vision software (Version 2.20, Iowa, USA)**.** COPD patients had a greater percentage of emphysema (%LAA_950_) compared to never-smoker and ever-smokers. * indicates, P < 0.05.

**Figure E2. (A) V**olcano plots showing the genes expressed specifically in the parenchyma, airways, and vessels of all patients who died of COVID-19. **B:** Boxplots showing the dynamic range of all genes assessed using the NanoString GeoMX DSP panel. **C:** Volcano plots showing the differential gene expression in the parenchyma, airways, and vessels of never- (NS) vs ever-smokers (ES) controls. For all volcano plots, the data were assessed using a generalized estimating equations (GEE) accounting for multiple ROIs per case, and was adjusted for age. The red dots indicate up-regulated genes and the blue dots down-regulated genes in COPD patients versus never- and ever-smokers for the specific tissue structures. The horizontal dotted line indicates the significance threshold of FDR<0.05, and the vertical dashed line indicates the effect size of 0. The grey dots indicate genes that were not significant at FDR<0.05. The top genes ranked by p-value are labelled with their gene symbols.

**Figure E3: (A)** Gene E (envelope) and (B) N (nucleocapside) RNA expression levels were measured in lungs from never- (NS) and ever- (ES) smoker controls and COPD patients who all died of SARS-CoV-2 infection using rt-PCR.

**Figure E4. (A)** Volcano plots showing the protein markers assessed in the parenchyma, airways, and vessels of all patients who died of COVID-19. The data were assessed using a generalized estimating equations (GEE) accounting for multiple ROIs per case, and was adjusted for age. The red dots indicate up-regulated genes and the blue dots down-regulated genes in COPD patients versus never- and ever-smokers for the specific tissue structures. The horizontal dotted line indicates the significance threshold of FDR<0.05, and the vertical dashed line indicates the effect size of 0. The grey dots indicate genes that were not significant at FDR<0.05. The top proteins ranked by p-value are labelled with their gene symbols. (**B**) Correlations between CD45RO protein expression and ACE2 protein expression in parenchyma, airways and vessels. The Rho and P values are given, p<0.05 was considered significant.

**Figure E5. (A)** CD45RO protein expression levels were quantified in the parenchyma, airways, and vessel regions of interest (ROIs) in lungs from patients with COPD versus ever- (ES) and never-smoker (NS) controls without SARs-CoV-2 infection, using GeoMX digital spatial profiling. The data were assessed using a generalized estimating equations (GEE) accounting for multiple ROIs per case, and was adjusted for age. The box plots represent the median and interquartile range, the error bars are the 5th and 95th percentile, and each dot represents a single ROI. * indicates P < 0.05, and *** indicates P < 0.001.
