## Supplementary figures and images for "Lung spatial profiling reveals a T cell signature in COPD patients with fatal SARS-CoV-2 infection"

### Supplemental Figure 1

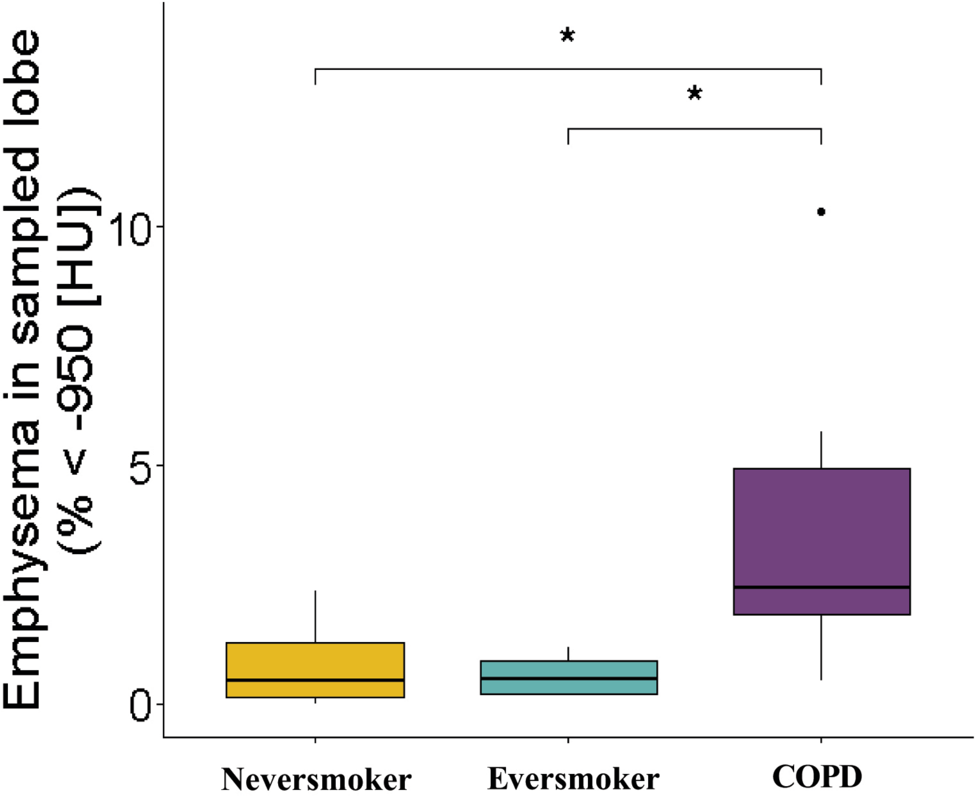

### Supplemental Figure 2

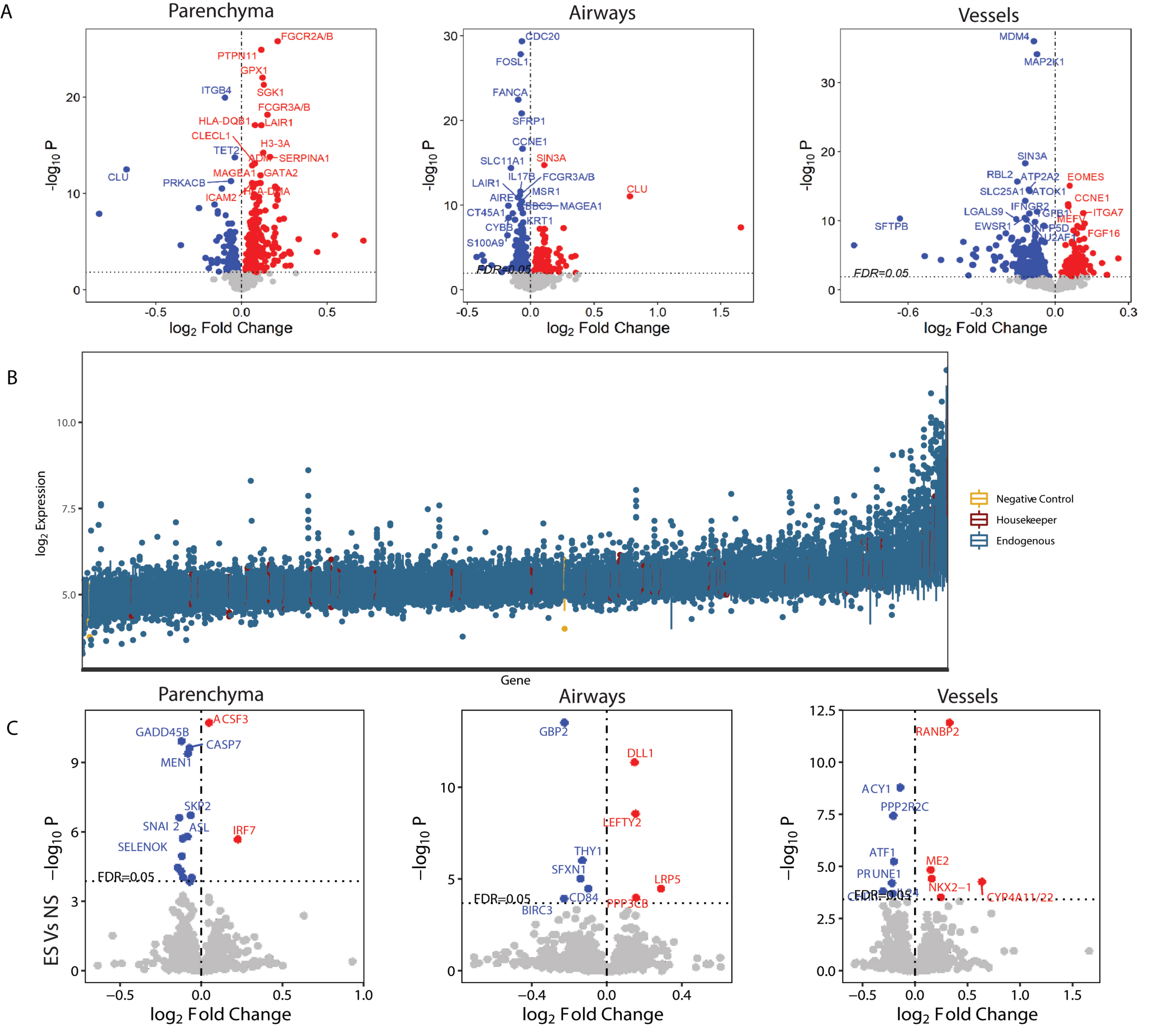

### Supplemental Figure 3

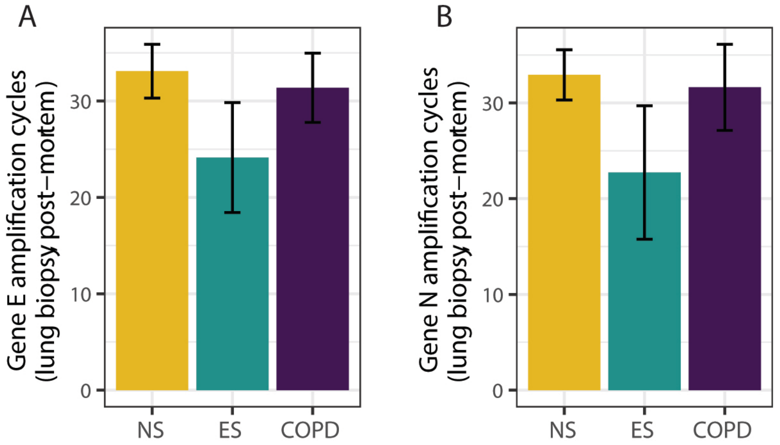

### Supplemental Figure 4

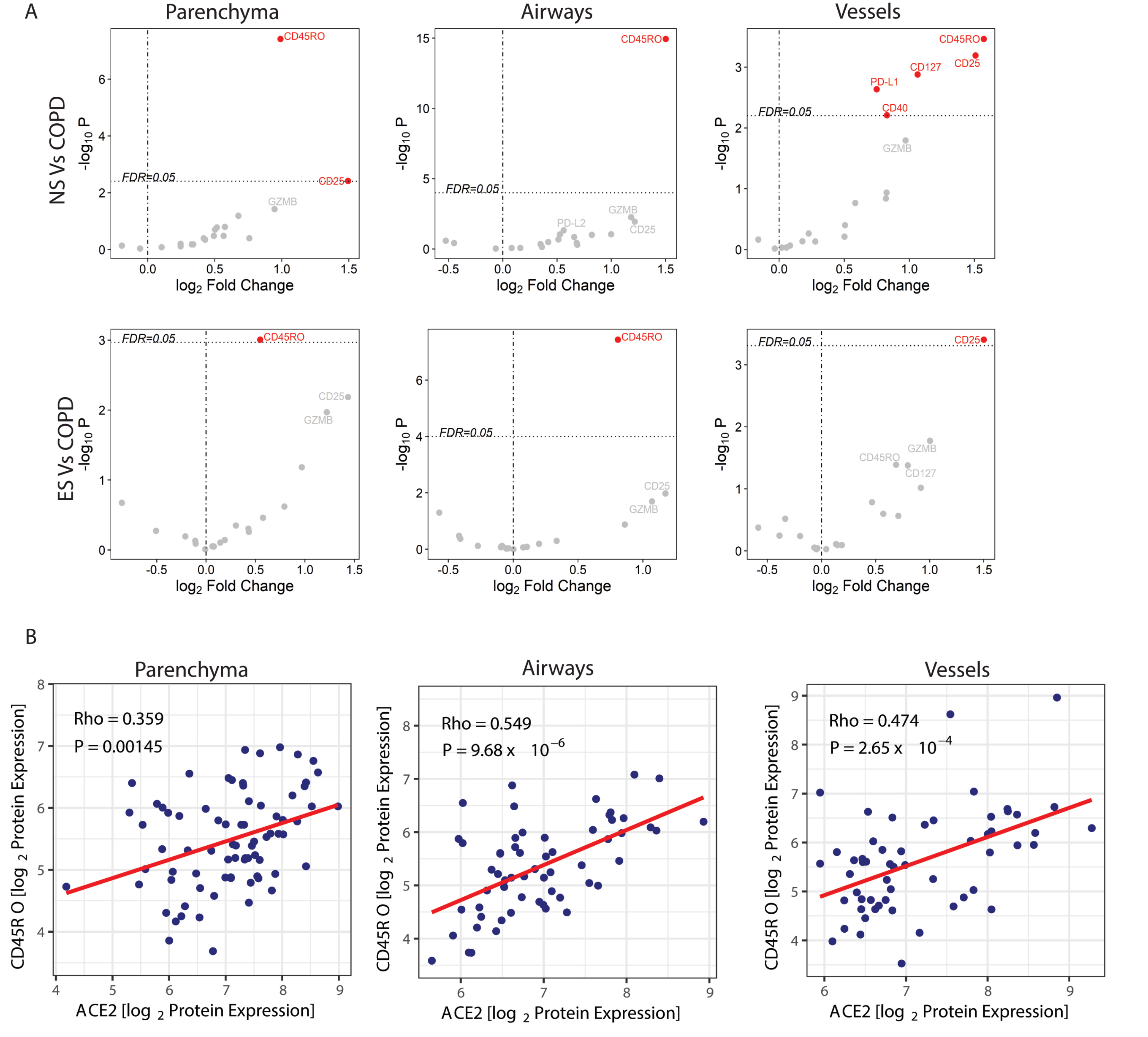

### Supplemental Figure 5

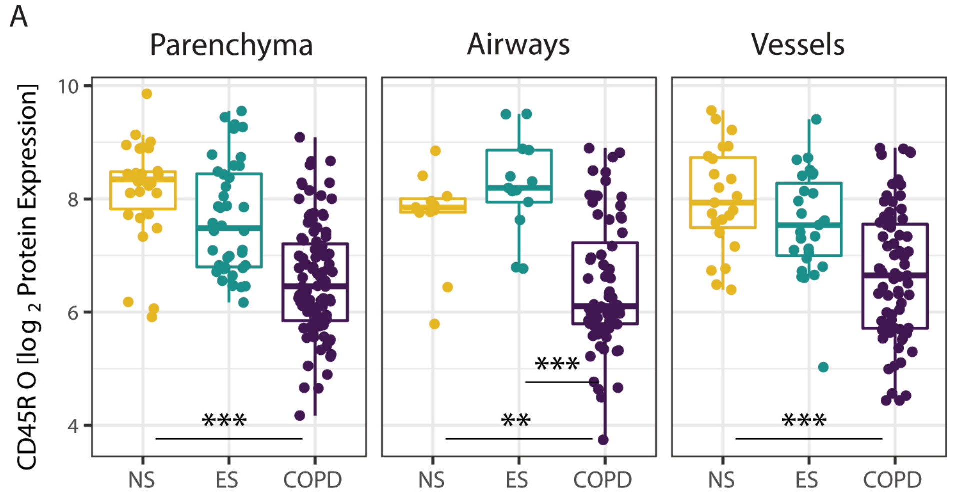
